# Bursting mitral cells time the oscillatory coupling between olfactory bulb and entorhinal networks in neonatal mice

**DOI:** 10.1101/2020.05.08.084079

**Authors:** Johanna K. Kostka, Sabine Gretenkord, Ileana L. Hanganu-Opatz

## Abstract

Shortly after birth, the olfactory system provides to blind, deaf, non-whisking and motorically-limited rodents not only the main source of environmental inputs, but also the drive boosting the functional entrainment of limbic circuits. However, the cellular substrate of this early communication remains largely unknown. Here we combine *in vivo* and *in vitro* patch-clamp and extracellular recordings to reveal the contribution of mitral cell (MC) firing to the early patterns of network activity in the neonatal olfactory bulb (OB) and lateral entorhinal cortex (LEC), the gatekeeper of limbic circuits. We show that MCs predominantly fire either in an irregular bursting or non-bursting pattern during discontinuous theta events in OB. However, the temporal spike-theta phase coupling is stronger for bursting MCs when compared to non-bursting cells. In line with the direct OB projections to LEC, both bursting and non-bursting firing augments during coordinated patterns of entorhinal activity, yet to a higher magnitude for bursting MCs. These cells are stronger temporally coupled to the discontinuous theta events in LEC. Thus, bursting MCs might drive the entrainment of OB-LEC network during neonatal development.

**KEY POINTS:**

- During early postnatal development mitral cells show either irregular bursting or non-bursting firing patterns
- Bursting mitral cells preferentially fire during theta bursts in the neonatal OB, being locked to the theta phase
- Bursting mitral cells preferentially fire during theta bursts in the neonatal lateral entorhinal cortex and are temporally related to both respiration rhythm- and theta phase
- Bursting mitral cells act as cellular substrate of the olfactory drive promoting the oscillatory entrainment of entorhinal networks

## INTRODUCTION

Cognitive performance mirrors complex interactions between limbic areas and is maximized throughout a permanent exchange with the environment. In rodents, these interactions emerge during early postnatal development with the hippocampus (HP) driving the functional assembling of prefrontal cortex (PFC), and LEC acting as a gatekeeper of both (Brockmann *et al.*, 2011; Hartung *et al.*, 2016; Ahlbeck *et al.*, 2018). However, the functional coupling of prefrontal-hippocampal-entorhinal circuits takes place during a developmental time window lacking most sensory inputs from the environment. Rodents are blind, deaf and do not whisker until the end of the second postnatal week. As a major exception, they are born with olfactory abilities (Welker, 1964). The functional olfactory system is not only critical for nursing and survival of pups (Teicher & Blass, 1977; Logan *et al.*, 2012) but seems to actively shape the functional coupling of limbic circuits (Gretenkord *et al.*, 2019). LEC receives direct input from OB, which, in contrast to other sensory systems, bypasses the thalamus (Luskin & Price, 1983; Igarashi *et al.*, 2012). As recently shown, coordinated patterns of electrical activity in the olfactory bulb that emerge either endogenously or odor-driven, gate the oscillatory entrainment of entorhinal networks at neonatal age (Gretenkord *et al.*, 2019). However, the cellular substrate of these early OB-LEC interactions is poorly understood.

MCs are the sole neuronal population of the OB projecting to the LEC. MCs of the adult OB have been extensively characterized in their passive and active properties (Balu *et al.*, 2004; Padmanabhan & Urban, 2010; Angelo & Margrie, 2011; Kollo *et al.*, 2014). According to the firing patterns, these neurons have been classified as regular firing and irregular bursting (or “stuttering”) (Yu *et al.*, 1993; Balu *et al.*, 2004; Nica *et al.*, 2010; Angelo & Margrie, 2011; Burton & Urban, 2014; Leng *et al.*, 2014). The MC firing patterns relate to the hyperpolarization-evoked sag potentials, irregular bursting MCs having less pronounced sag currents when compared to regular firing MCs (Angelo & Margrie, 2011; Burton & Urban, 2014). Moreover, the sag currents have been identified as a biophysical fingerprint of the glomerular network affiliation and sensory processing (Angelo *et al.*, 2012). MCs are part of a complex neuronal circuitry that enables odor information coding (Wouterlood *et al.*, 1985; Adam & Mizrahi, 2011). Their axons project onto apical dendrites of pyramidal and stellate cells in layer II/III of LEC, which in turn give rise to the perforant path projections to the hippocampal formation (Witter, 2007).

The MC projections to LEC emerge already at embryonic stage (Walz *et al.*, 2006; Hirata *et al.*, 2019) and are fully developed at the postnatal age when prefrontal-hippocampal-entorhinal circuits generate coordinated patterns of oscillatory activity. While the morphology and connectivity of MCs across development have been analyzed in detail (Hinds & Hinds, 1976; Lin *et al.*, 2000; Blanchart *et al.*, 2006), few *in vitro* data documented their biophysical properties at early ages (Yu *et al.*, 2015). A slice preparation might be instrumental for linking MC properties to a very local circuit within the OB (Angelo *et al.*, 2012), yet does not allow to identify their contribution to the long-range communication within OB-LEC circuits.

For addressing this aim, we combined *in vivo* patch-clamp recordings with extracellular recordings of local field potential (LFP) and spiking activity in the OB and LEC of urethane-anesthetized neonatal (postnatal day [P] 8-10) mice. We show that, similar to *in vitro* conditions, MCs show two different firing patterns *in vivo*, non-bursting and irregular bursting, the latter one being stronger temporally correlated with the discontinuous theta bursts in the OB. Moreover, the irregular bursting MCs time the oscillatory rhythms in the LEC. These data identify the irregular bursting subpopulation of MCs as the main drivers of long-range coupling between OB and limbic circuitry.

## METHODS

### Ethical Approval

All experiments were performed in compliance with the German laws and the guidelines of the European Union for the use of animals in research (European Union Directive 2010/63/EU) and were approved by the local ethical committee (Behörde für Gesundheit und Verbraucherschutz Hamburg, ID 15/17).

### Animals

Time-pregnant C57Bl/6/J mice from the animal facility of the University Medical Center Hamburg-Eppendorf were housed individually in breeding cages at a 12h light / 12h dark cycle and fed ad libitum. Male mice underwent either *in vitro* or *in vivo* whole-cell patch clamp or *in vivo* multi-side electrophysiological recordings at P8–10 using experimental protocols similar to those described previously (Bitzenhofer *et al.*, 2015; Bitzenhofer *et al.*, 2017; Gretenkord *et al.*, 2019). In line with the 3R (reduce, refine, replace) principles of animal protection, part of the data resulted from mice that have been investigated for a different data set in a previous publication (Gretenkord *et al.*, 2019).

### Surgical procedures and recordings

#### Surgical procedure for electrophysiology *in vitro*

For patch-clamp recordings, pups were decapitated, and brains were sliced with a vibratome (Leica, VT1000S) into 300 μm thick coronal sections in ice-cooled oxygenated high-sucrose-based artificial cerebral spinal fluid (ACSF) containing (in mM): 228 sucrose, 2.5 KCl, 1 NaH_2_PO_4_,1 26.2 NaHCO_3_, 11 glucose, 7 MgSO_4_ (310 mOsm). Prior to recordings, slices were maintained at room temperature and superfused with oxygenated ACSF containing (in mM): 119 NaCl, 2.5 KCl, 1 NaH_2_PO_4_, 26.2 NaHCO_3_, 11 glucose, 1.3 MgSO_4_ (310 mOsm) at 37 °C.

### *In vitro* whole-cell patch-clamp recordings

Whole-cell patch-clamp recordings were performed from neurons identified by their location in the MC layer and their cell body size. All recordings were performed at room temperature. Recording electrodes (4 – 9 MΩ) were filled with K-gluconate based solution containing (in mM): 130 K-gluconate, 10 HEPES, 0.5 EGTA, 4 Mg-ATP, 0.3 Na-GTP, 8 NaCl (285 mOsm [pH 7.4]), and 0.5% biocytin for post hoc morphological identification of recorded cells. Recordings were controlled with the Ephus software (Suter *et al.*, 2010) in the MATLAB environment (The MathWorks, MA, USA). Capacitance artefacts were minimized using the built-in circuitry of the patch-clamp amplifier (Axopatch 200B; Molecular devices, CA, USA). The signals were low-pass filtered at 10 kHz and recorded online. All potentials were corrected for the liquid junction potential of the gluconate-based electrode solution, which according to our measurement was -8.65 mV. The resting membrane potential (RMP) was measured immediately after obtaining the whole-cell configuration. For the determination of input resistance (R_in_), membrane time constant (τ_m_) and membrane capacitance (C_m_) hyperpolarizing current pulses (−60 pA) of 600 ms duration were applied from resting membrane potential. Analysis was performed offline using custom-written scripts in the MATLAB environment (MathWorks, Natick, MA).

### Surgical procedure for electrophysiology *in vivo*

Mice were injected i.p. with urethane (1 mg/g body weight; Sigma-Aldrich, MO, USA) prior to surgery. Under isoflurane anesthesia (induction: 5%, maintenance: 2.5%) the head of the pup was fixed into a stereotaxic apparatus using two plastic bars mounted on the nasal and occipital bones with dental cement. The bone above the right OB (0-5-0.8 mm anterior to fronto-nasal suture, 0.5 mm lateral to inter-nasal suture) as well as LEC (0 mm posterior to lambda, 6-6.5 mm lateral from the midline) was carefully removed by drilling a hole of <0.5 mm in diameter. Throughout surgery and recording session the mice were maintained on a heating blanket at 37°C.

### Multi-site electrophysiological recordings *in vivo*

One-shank electrodes (NeuroNexus, MI, USA) with 16 recording sites (0.4-0.8 MΩ impedance, 50 µm inter-site spacing for OB, 100 µm inter-site spacing for LEC) were inserted into OB (0.5 - 1.8 mm, angle 0°) as well as in LEC (depth: 2 mm, angle: 10° from the vertical plan). Before insertion, the electrodes were covered with DiI (1,1’-Dioctadecyl-3,3,3’,3’-tetramethylindocarbocyanine perchlorate, Molecular Probes, Eugene, OR). A silver wire was inserted into the cerebellum and served as ground and reference electrode. Before data acquisition, a recovery period of 20 min following the insertion of electrodes was provided. Extracellular signals were band-pass filtered (0.1 Hz – 9 kHz) and digitized (32 kHz or 32,556 kHz) by a multichannel amplifier (Digital Lynx SX; Neuralynx, Bozeman, MO; USA) and Cheetah acquisition software (Neuralynx). Spontaneous activity was recorded for 30 min before insertion of glass electrode for patch-clamp recordings in OB. The position of recording electrodes in OB and LEC was confirmed after histological assessment *post-mortem*. For the analysis of LFP in OB, the recording site centered in external plexiform layer was used, whereas for the analysis of spiking activity all recording sites located in the MC layer were considered. For analysis of LFP in LEC, only recording sites that were histologically confirmed to be located in superficial entorhinal layers were used.

### *In vivo* whole-cell patch-clamp recordings

Whole-cell patch-clamp recordings were performed using glass pipettes (4 - 9 MΩ) that were filled with (in mM) 135 K-gluconate, 4 KCl, 10 Na2-phosphocreatine, 10 HEPES, 4 MgATP, 0.3 NaGTP for current clamp recordings. In all experiments 0.5% biocytin (Sigma-Aldrich) was included in the pipette solution for later morphological identification of the recorded cells. Patch pipettes were inserted over the OB and advanced at 35° from the vertical plane into the brain until a depth of 150 µm. To avoid clotting of the pipette, positive pressure (250 mbar) was applied to the pipette when advancing it through the dorsal OB. In the proximity of the LFP electrode, the pressure was reduced and the pipette slower advanced (1μm/s). When the pipette blindly contacted a cell, the current amplitude in response to a 10 mV-test pulse decreased. Action potentials were recorded using a discontinuous voltage-clamp/current-clamp amplifier (ELC-03XS, npi elektronik, Tamm, Germany). The signals were amplified and low-pass filtered at 3 kHz, visualized on an oscilloscope (Tektronix, Beaverton, OR), digitized online with an AD/DA board, recorded and processed with Cheetah software. All potentials were corrected for the liquid junction potential, which according to our measurement was -10.37 mV. Similar to *in vitro* recordings resting membrane potential (RMP) was measured immediately after obtaining the whole-cell configuration. For the determination of input resistance (R_in_), membrane time constant (τ_m_) and membrane capacitance (C_m_) hyperpolarizing current pulses (−60 pA) of 500 ms duration were applied from resting membrane potential. Recording traces were analyzed offline using custom-written scripts in the MATLAB environment (MathWorks, Natick, MA).

### Morphology

Mice were anesthetized with 10% ketamine (aniMedica, Germany) / 2% xylazine (WDT, Germany) in 0.9% NaCl solution (10 µg/g body weight, i.p.) and transcardially perfused with Histofix (Carl Roth, Germany) containing 4% paraformaldehyde (PFA). Brains were postfixed in 4% paraformaldehyde for 24 h. Tissue blocks containing the OB were sectioned in the coronal plane at 400 μm using a vibratome. 300 µm thick slices from *in vitro* recordings were postfixed in 4% paraformaldehyde for 24 h. After fixation brain slices were washed in PBS, blocked and permeabilized (0.8% Triton, Sigma-Aldrich, 5,0% normal bovine serum albumin, Jackson Immuno Research, 0.05% sodium azide, Sigma-Aldrich). Staining with Cyanine dye 2 (Cy2)-conjugated streptavividin (Jackson Immuno Research, 1:400 in PBS, with 3% bovine serum albumin, Jackson Immuno Research and 0.05% sodium azide, Sigma-Aldrich) for 60 min was used to identify biocytin-filled cells. After washing in PBS, slices where mounted with Fluoromount (Sigma-Aldrich). The positions of the DiI-labeled extracellular electrodes in the OB and LEC were reconstructed using fluorescence microscopy. The morphology of recorded neurons was examined in more detail using confocal microcopy (DM IRBE, Leica, Germany). Image stacks along the z-axis were made from brain slices using an argon laser to excite the fluorophore Cy2 at a wavelength of 488 nm, while the emitted light was detected at around 510 nm.

### Data Analysis

#### LFP analysis

Data were analyzed offline using custom-written scripts in the MATLAB environment (MathWorks, Natick, MA). Data were first low-passed filtered (<100 Hz) using a third-order Butterworth filter before down-sampling by factor 20 to 1.6 kHz or 1.65 kHz to analyze LFP. All filtering procedures were performed in a manner preserving phase information.

#### Detection of oscillatory activity

Discontinuous network oscillations in the LFP recorded from OB and LEC were detected using a previously developed unsupervised algorithm (Cichon *et al.*, 2014). Briefly, deflections of the root mean square of band-pass filtered (1-100 Hz) signals exceeding a variance-depending threshold (2.5 times the standard-deviation from the mean) were assigned as oscillatory periods. Only oscillatory periods lasting at least 1 s were considered for analysis.

#### Power spectral density

Power spectral density was analyzed for either the entire signal or 3 s long periods with theta oscillations, which were concatenated and the power was calculated using Welch’s method with non-overlapping windows. Relative LFP power was calculated by dividing the power of periods with theta oscillations by the power of periods without theta oscillations. Time-frequency plots of power were calculated with a continuous wavelet transform (Morlet wavelet).

#### Single unit clustering

Single units were automatically detected and clustered using the python-based software klusta (Rossant *et al.*, 2016) and manually curated using phy (https://github.com/cortex-lab/phy).

#### Analysis of the passive membrane properties

For all intracellular recorded neurons RMP, input resistance (R_i_), membrane time constant (t), membrane capacity (C_m_), action potential (AP) amplitude, halfwidth and firing threshold where calculated. The Input resistance (R_i_) was calculated according to the Ohm’s law by dividing the resulting potential changes by the amplitude of the injected current (−60 pA) (Barbour, 2011). The membrane time constant (τ) was calculated by fitting a monoexponential function to the induced potential deflection. Membrane capacity (C_m)_ was calculated by dividing the membrane time constant by the membrane resistance. Firing threshold was calculated by estimating the voltage with the steepest increase in slope of the phase plot (dV/dt against V). Sag amplitude was calculated for each cell as the difference between the initial voltage response and the steady state response to a hyperpolarizing current pulse of -100 pA. For this, the initial voltage response was measured from the minimal voltage reached in the first 200 ms of the current pulse. The steady-state voltage response was estimated by taking the mean value of the last 100 ms of the voltage response.

#### Analysis of action potential firing properties

Action potential frequency, inter-spike intervals (ISI) and coefficient of variation (CV) were calculated for all recorded units with firing rates over 0.1 Hz. The firing rate was calculated by dividing the number of spikes during the whole signal or during theta events by the duration of the analyzed periods. Action potential firing during theta burst and non-burst periods were normalized by the total number of spikes of the corresponding unit. Mean action potential firing during theta oscillations was estimated by aligning the spikes of one unit to the start of each theta burst and using a Gaussian kernel function (delta = 0.1) (Shinomoto, 2010). CV was calculated by dividing the standard deviation of the ISI of each neuron by its mean ISI.

#### Burst detection

Burst detection of spontaneous action potential (AP) firing was achieved as previously described (Gorin *et al.*, 2016). APs where considered in a burst if at least 4 consecutive APs had smaller Inter-spike-intervals as the median (ISI) calculated from the corresponding Poisson distribution. Neurons where considered as bursting if at least 50% of their APs occurred in bursts. Autocorrelation histograms where calculated for time windows of 120 s-length.

### Assessment of relationships between LFP and spiking activity

#### Spike-LFP coupling

Phase locking of spiking units to network oscillations was assessed using a previously described algorithm (Siapas *et al.*, 2005). For this, the LFP signal was bandpass filtered (2-4 Hz (RR), 4-12 Hz (theta)) using a third-order Butterworth filter. The instantaneous phase was extracted using the Hilbert transform on the filtered signal. The coupling between spikes and network oscillations was tested for significance using the Rayleigh test for non-uniformity. Only neurons that showed significant phase locking were considered for the analysis of locking strength which was calculated as mean resulting vector length.

#### Pairwise phase consistency

Pairwise phase consistency (PPC) was computed as previously described (Vinck *et al.*, 2010). For this, the phase in the band of interest was extracted using the Hilbert transform of the band-pass filtered LFP trace. The mean of the cosine of the absolute angular distance (dot product) among all pairs of phases was calculated.

#### Spike-triggered average

Spike-triggered average was calculated by taking the mean of the band-pass filtered LFP (2-4 Hz (RR), 4-12 Hz (theta)) for 1 s-long time-windows centred on each spike.

### Statistics

Statistical analysis was performed in MATLAB environment. Gaussian distribution of the data was assessed using the Kolmogorov-Smirnov test for sample size >10. None of the data sets were normally distributed. Therefore, data were tested for significance using Wilcoxon rank-sum test (2 unrelated samples) or Wilcoxon sign-rank test (2 related samples). Values were considered as outliers and removed when their distance from the 25^th^ or 75^th^ percentile exceeded 1.5 times the interquartile interval. Phase locking was tested for significance using the Rayleigh test for non-uniformity. Phase data where compared using a circular, non-parametric circular test similar to the Kruskal-Wallis test (*circ_cmtest* function).

## RESULTS

### Passive and active membrane properties of neonatal mitral cells

We firstly characterized the membrane properties of mitral cells (MCs) at neonatal age. For this, we performed whole-cell patch-clamp recordings from MCs in the OB of P8-10 mice *in vivo* (n=12) and *in vitro* (n=11 cells) (Figure 1A). MCs showed similar passive membrane properties under *in vivo* and *in vitr*o conditions (Table 1). Solely the membrane time constant and input resistance where slightly higher for the *in vitro* recorded cells when compared to those recorded under *in vivo* conditions (Table 1). Similar differences between the two conditions have been described for pyramidal neurons in the visual cortex of adult rats (Monier *et al.*, 2008). Independent of the recording conditions, all MCs showed almost linear voltage-current relationships and increased their firing in response to depolarizing current injection (Figure 1C & D). MCs recorded under *in vitro* and *in vivo* conditions did not differ in their active membrane properties (i.e. action potential threshold, spontaneous firing rate (Table 1)).

**Table 1:** Passive and active membrane properties of neonatal MCs recorded intracellular *in vivo* and *in vitro*.

|  | *in vivo* intracellular<br>(median, [iqr]) | *In vitro* intracellular<br>(median, [iqr]) | p<br>(Wilcoxon<br>rank-sum<br>test) |
| --- | --- | --- | --- |
| **RMP (mV)** | -59 [-63 – -56.5] | -58.31 [-62.19 – -55.7] | 0.601 |
| **C<sub>m</sub> (pF)** | 54.22 [33.74 – 71.40] | 70.41 [46.51 – 95.65] | 0.340 |
| **R<sub>i</sub> (MΩ)** | 330.46 [247.25 – 406.95] | 550.32 [399.32 – 689.92] | 0.018 |
| **τ<sub>m</sub> (ms)** | 18.57 [11.94 – 22.84] | 33.34 [19.20 – 36.16] | 0.010 |
| **Log firing rate (Hz)** | -1.78 [-2.43 – -0.34] | 0.44 [-0.07 – 0.70] | 0.055 |
| **AP threshold (mV)** | -52.46 [-55.77 – -45.91] | -50.10 [-52.58 – -48.61] | 0.815 |

**Figure 1.**
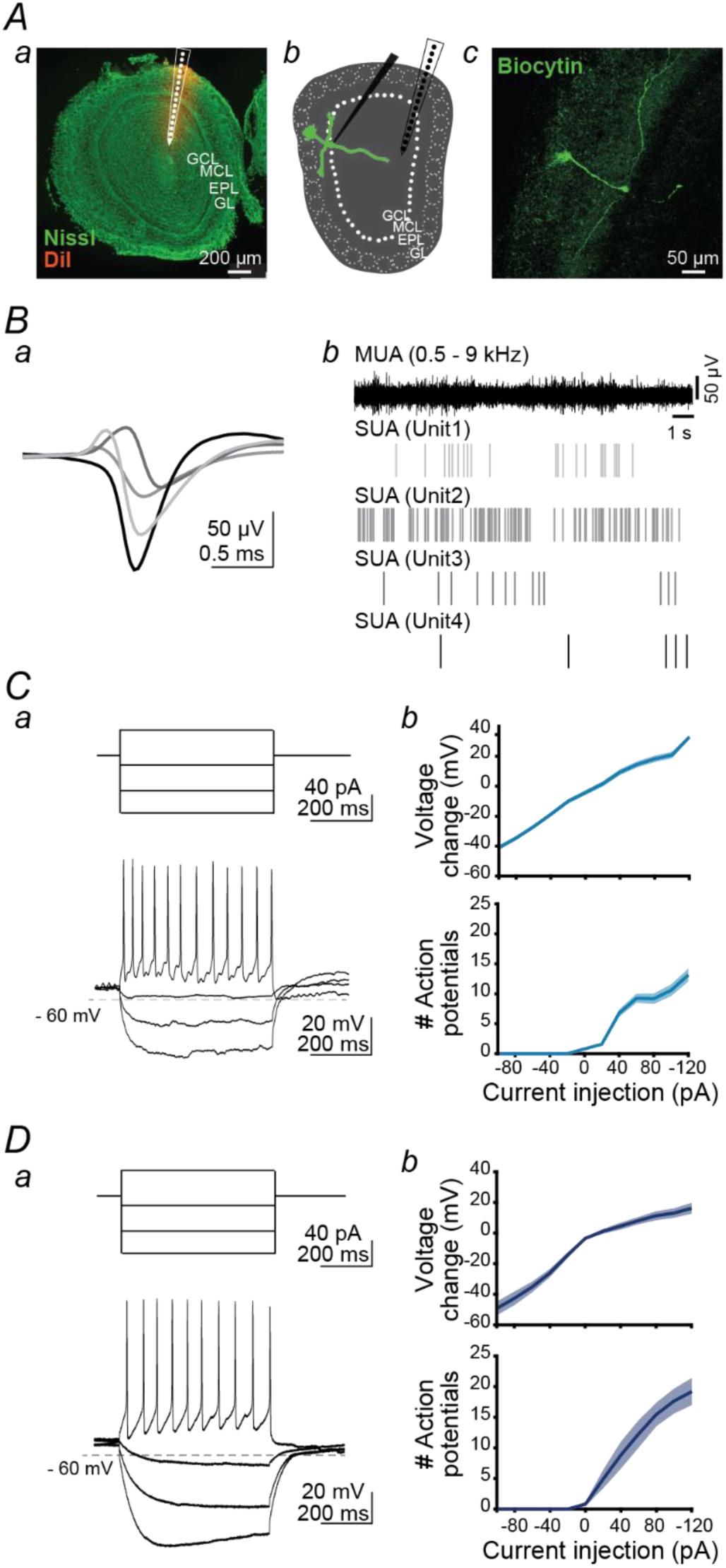
Passive and active membrane properties of neonatal mitral cells *in vivo* and *in vitro*. A, Digital photomontage reconstructing the track of the multi-site DiI-labeled recording electrode (red) in a Nissl-stained coronal section of the OB from a P9 mouse (a). Schematic showing the recording configuration for simultaneous LFP and whole-cell patch-clamp recordings in the dorsal OB. GCL= granule cell layer, MCL= mitral cell layer, EPL: external plexiform layer, GL: glomerular layer (b). Confocal image displaying a biocytin-filled mitral cell (green) in the OB of a P8 mouse (c). B, Example of single units with distinct waveforms recorded in the MC layer of the OB from a P8 mouse (a). Extracellular recording of the MUA from the OB displayed together with the raster plots of the four units shown in (a). C, Current-voltage response of a MC recorded in the OB of a P9 mouse *in vivo* (a). I/V relationship and firing rate in relationship to current averaged for all MC recorded *in vivo* (b). D, Same as in C for MCs recorded in slice preparation.

These data indicate that in slice preparation the MCs maintain their membrane properties detected in the OB of neonatal mice *in vivo*.

### Neonatal mitral cells fire either busting or non-bursting

Next, we investigated the spontaneous firing patterns of MCs at neonatal age *in vivo* and *in vitro*. Extracellular recording of multiunit activity (MUA) in the OB of anesthetized P8-10 mice using multi-site silicon probes (Figure 1A(a)) was performed in addition to whole-cell patch-clamp recordings. MUA was clustered for identification of single units (SUA) using offline analysis tools (Rossant *et al.*, 2016) (n=78 cells) (Figure 1B). In order to characterize spontaneous firing patterns of MCs, we analyzed the timing of firing using a previously described method (Gorin *et al.*, 2016). Action potentials (APs) were considered to be part of a burst if at least 4 consecutive APs had smaller inter-spike intervals (ISI) than the median ISI calculated from the corresponding Poisson distribution. MCs were classified as bursting if at least 50% of their APs occurred in bursts and their ISI distribution was significantly different from the expected Poisson distribution (Figure 2A(c) & B(c)). Moreover, bursting MCs were identified by the presents of several peaks in the autocorrelation histograms (Figure 2A(b) & B(b)). We analyzed the firing pattern of MCs when (i) extracellularly recorded as units *in vivo*, (ii) intracellularly patch-clamp recorded *in vivo*, and (iii) intracellularly patch-clamp recorded *in vitro*. In all three recording conditions bursting and non-bursting MCs were detected (Figure 2A & B). Moreover, the proportion of MCs classified as bursting was comparable in the two recording conditions *in vivo* (Figure 2C; extracellular: 66.2 %, intracellular: 66.7 %), whereas fewer bursting neurons were detected among the *in vitro* recorded MCs (57.1 %). This difference might result from the reduced connectivity and inputs of MCs in a slice preparation.

**Figure 2.**
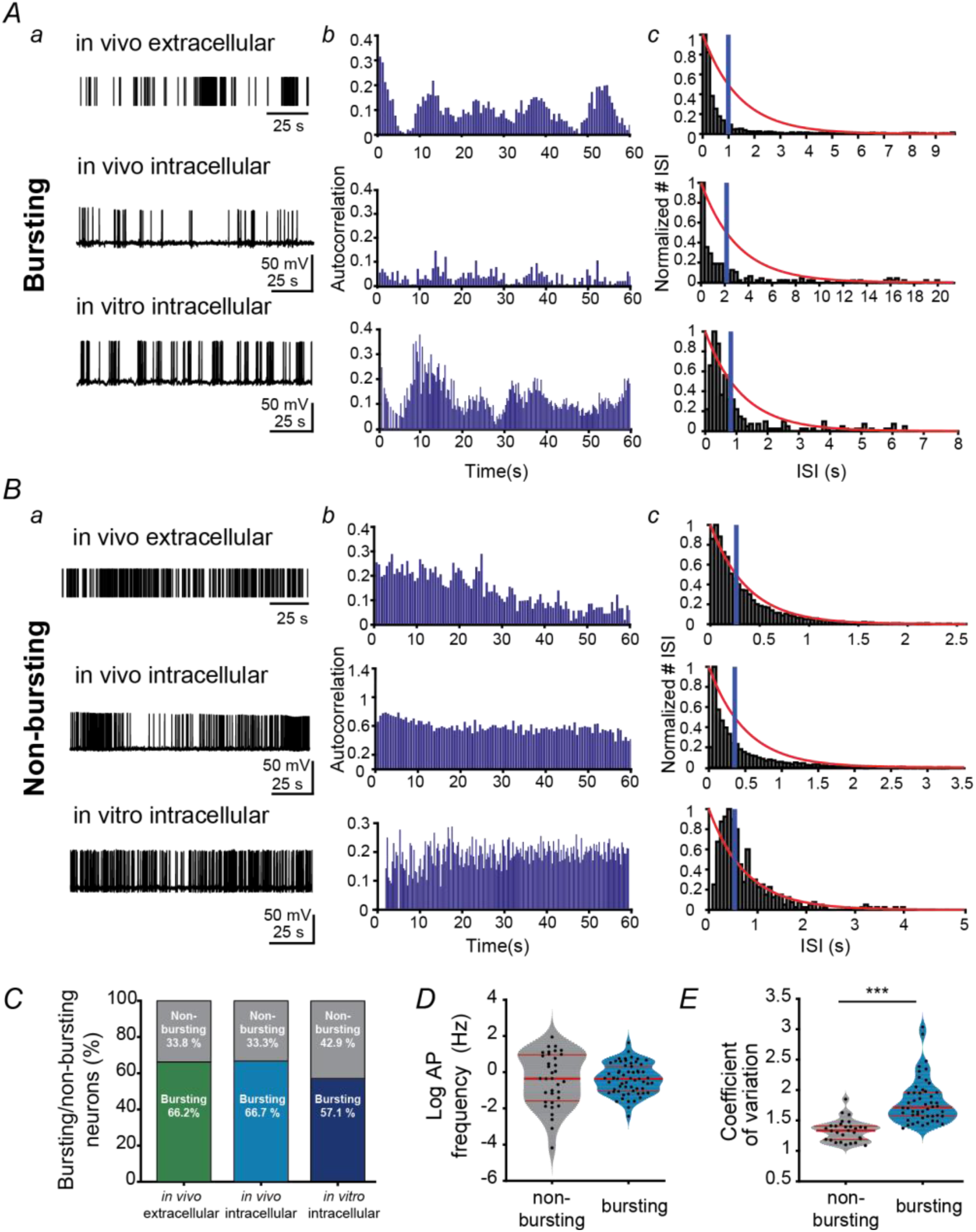
Irregular bursting and non-bursting mitral cells are present in the neonatal OB. A, Example of bursting MCs extracellularly recorded *in vivo* (top), patch-clamp recorded *in vivo* (middle) and patch-clamp recorded *in vitro* (bottom) (a). Autocorrelation (b) and inter-spike interval (ISI) histograms (c) displayed for each neuron shown in (a). The blue line corresponds to median ISI. The probability curves of a Poisson process with the same firing rate are displayed in red. B, Same as B for non-bursting MCs. C, Bar plot displaying the percentage of bursting vs. non-bursting MCs for each recording condition. D, Violin plots showing the log transformed firing frequency for bursting vs. non-bursting MCs. E, Violin plots showing the coefficient of variation of ISIs for bursting vs. non-bursting MCs. (*** p < 0.001, Wilcoxon rank-sum test). For violin plots the red lines correspond to median (thick line) as well as 1. and 3. quantiles of the distribution (thin lines).

The firing rates of bursting and non-bursting MCs were similar and therefore, not instrumental for the classification of cells (Figure 2D, Table 2). Moreover, the passive membrane properties of intracellularly recorded MCs did not differ between the two groups (Table 2). In contrast, the coefficient of variation (CV) of inter-spike intervals (ISI), a measure of spike irregularity (Figure 2E, Table 2), was higher for bursting MCs. Visual examination and CV > 1 revealed that all MCs fire irregularly, but the degree of irregularity was higher in bursting MCs.

**Table 2:** Passive and active membrane properties of neonatal non-bursting and bursting MCs.

|  | Non-bursting MCs<br>(median, [iqr]) | Bursting MCs<br>(median, [iqr]) | p<br>(Wilcoxon<br>rank-sum<br>test) |
| --- | --- | --- | --- |
| RMP (mV) | -58.03 [-66.46 – -50.45] | -58.31 [-59.25 – -56.52] | 0.951 |
| C<sub>m</sub> (pF) | 72.97 [42.84 – 84.58] | 53.71 [41.31 – 62.79] | 0.167 |
| R<sub>i</sub> (MΩ) | 569.12 [257.73 – 723.46] | 362.48 [279.19 – 427.68] | 0.156 |
| τ<sub>m</sub> (ms) | 16.20 [9.67 – 19.41] | 22.82 [14.56 – 29.80] | 0.083 |
| Log firing rate (Hz) | -0.35 [-1.59 – 0.96] | -0.37 [-1.04 – 0.32] | 0.885 |
| CV | 1.33 [1.19 – 1.41] | 1.72 [1.57 – 1.96] | 1.9 *10<sup>-9</sup> |

Previous studies proposed a causal link between the presence of a hyperpolarization-evoked sag potential that are mediated by hyperpolarization-activated cyclic nucleotide gated (HCN) channels, and the firing patterns of MCs (Angelo & Margrie, 2011; Burton & Urban, 2014; Yu *et al.*, 2015). To uncover whether this link exists already during development, we analyzed the sag potentials in MC *in vivo* and *in vitro* (Figure 3A). While both bursting and non-bursting MCs showed such hyperpolarization-evoked potentials, their amplitude is larger in non-bursting MCs (Figure 3B & C; non-bursting (nB): median: 12.805 mV, iqr: 4.124 – 26.766 mV; bursting (B): median: 3.302, iqr: 0.062 – 9.165 mV; p = 0.043, Wilcoxon rank-sum test). These data are in line with the previously reported properties of adult MCs, among which bursting (stuttering) MCs have smaller sag potentials (Angelo & Margrie, 2011; Burton & Urban, 2014; Yu *et al.*, 2015).

**Figure 3.**
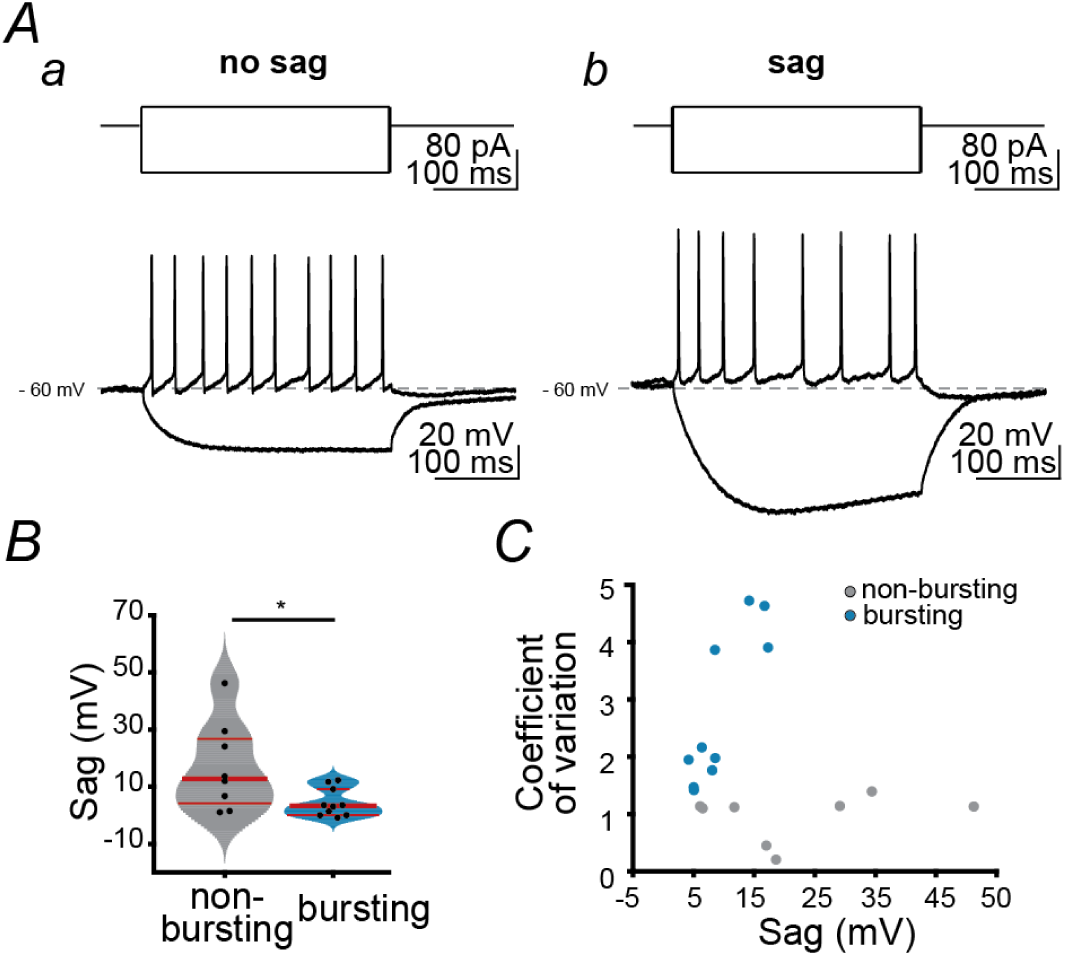
Sag current in neonatal bursting and non-bursting mitral cells. A, Voltage responses of MCs without (a) and with sag current (b) to the injection of hyper- and depolarizing current pulses at resting membrane potential. A suprathreshold current pulse elicits repetitive action potentials. B, Violin plot displaying the magnitude of sag potentials for bursting vs. non-bursting MCs. (*p < 0.05, Wilcoxon rank-sum test). Red lines correspond to median (thick line) as well as 1. and 3. quantiles of the distribution (thin lines). C, Dot plot displaying the coefficient of variation in relationship to the magnitude of sag potential for all investigated non-bursting (gray) and bursting (blue) MCs.

Taken together, these findings indicate that bursting and non-bursting MCs represent two distinct subpopulations which most likely differ in their expression of HCN channels.

### Mitral cells preferentially fire during theta events in the OB

The presence of two populations of MCs with different firing patterns raises the questions whether they differently relate to the network activity in OB. In a previous study, we showed that MCs critically contribute to the oscillatory entrainment of OB in neonatal mice (Gretenkord *et al.*, 2019). Shortly after birth the OB shows two patterns of coordinated activity, a continuous respiration-related rhythm (RR) with peak frequencies of 2-4 Hz and discontinuous theta events with frequencies within 4-12 Hz range (Figure 4A & B). To assess the relationship between individual MCs and network oscillations, we used two distinct approaches. First, we monitored the temporal relationship between the MCs activity and the simultaneously recorded LFP in the OB of P8-10 mice. Bursting and non-bursting neonatal MCs showed significant higher normalized firing rates during theta events (nB: median: 1.320, iqr: 1.159 - 1.450; B: 1.623, iqr: 1.461 - 1.766) when compared to periods without theta events (nB: median: 0.760, iqr: 0.630 – 0.882; B: median: 0.551, iqr: 0.453 – 0.667, p(nB) = 9.78*10^−6^, p(B) = 3.5*10^−10^, Wilcoxon sign-rank test) (Figure 4A, C & D). However, the two MC populations are not identically entrained by the theta events, since bursting MCs fire stronger during these events than non-bursting MCs (Figure 4D; p = 7.80*10^−7^, Wilcoxon rank-sum test). In contrast, in the absence of theta events, the non-bursting MCs have higher normalized firing rates when compared to bursting MCs (p = 5.19*^-7^, Wilcoxon rank-sum test). These data provide the first evidence that bursting MCs preferentially fire during theta events.

**Figure 4.**
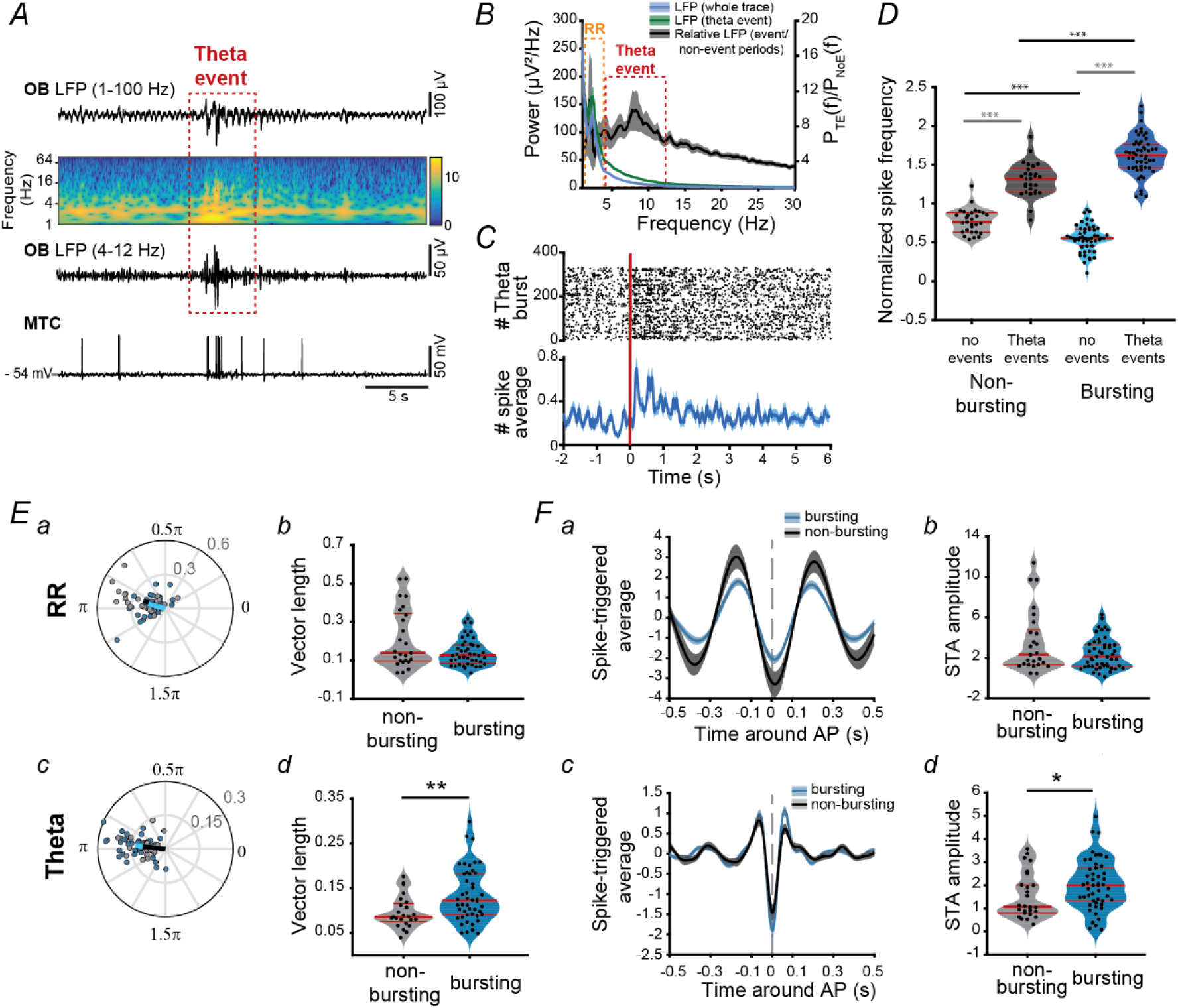
Bursting mitral cells fire preferentially during OB theta events. A, Top, extracellular LFP recording of the oscillatory activity in the OB of a P9 mouse displayed after band-pass filtering (1-100 Hz, 4-12 Hz) and accompanied by the wavelet spectrogram at identical timescale. Bottom, action potentials recorded simultaneously with the LFP from a MC at resting membrane potential in whole-cell current-clamp configuration. Note the temporal correlation between the discontinuous theta event (marked by red box) and action potentials. B, Power spectra (mean ± SEM) of LFP in OB for whole recording (blue), during discontinuous bursts (green) as well as of theta bursts normalized to non-bursting activity (black). C, Representative raster plot (top) and averaged number of spikes (bottom) from an extracellularly recorded bursting unit aligned to the start time of each theta event (red line). D, Violin plot displaying the normalized firing rates of *in vivo* recorded non-bursting (gray) and bursting (blue) MCs during theta events (dark colors) and time windows lacking those (light colors). E, Phase locking of MC units to RR and theta events in OB. Polar plots displaying the phase locking of significantly locked bursting (blue) and non-bursting (gray) units to RR (a) and theta (c). Violin plots showing the mean resulting vector length for bursting (blue) and non-bursting units phase locked to RR (b) and theta (d). Only significantly locked units where used for analysis. F, Spike triggered average for RR (a) and theta (c) around the time of action potential occurrence for bursting (blue) and non-bursting (gray) MCs. Corresponding violin plots showing the absolute STA amplitude around the time of action potential occurrence for RR (b) and theta (d). (*p<0.05, **p<0.01, *** p < 0.001; Wilcoxon rank-sum test (black) or Wilcoxon sign-rank test (gray)). For violin plots the red lines correspond to median (thick line) as well as 1. and 3. quantiles of the distribution (thin lines).

Second, we investigated whether OB network oscillations time the firing of single units extracellularly recorded in the MC layer of P8-10 mice. For this, we calculated the phase locking of spikes from putatively MCs to RR and theta events, respectively. A high proportion of non-bursting and bursting MCs where locked to RR (non-bursting: 96 %, bursting 88%) as well as to the theta rhythm (non-bursting: 96%, bursting 94%). However, the locking strength of significantly locked bursting MCs estimated by the mean resulting vector length to the theta rhythm was higher than of non-bursting MCs (Figure 4E(d); nB: median: 0.086. iqr: 0.075 – 0.116 B: median: 0.122, iqr: 0.091 – 0.182; p = 0.005; Wilcoxon rank-sum test). In contrast, the locking strength of both MC populations to RR was similar (Figure 4E(b); nB: median: 0.140, iqr: 0.097 – 0.342; B: median: 0.126, iqr: 0.084 – 0.181; p = 0.166, Wilcoxon rank-sum test). Taking into account that phase locking strongly depends on the firing rate, we confirmed the results above by calculating the pair-wise phase consistency (PPC) (Vinck *et al.*, 2010). While both MC populations had similar PPC for RR (nB: median: 0.015, iqr: 0.008 – 0.080, B: median: 0.012, iqr: 0.004 – 0.025; p = 0.160; Wilcoxon rank-sum test), for the coupling with theta events the PPC values were larger for bursting MCs when compared to the values for non-bursting MCs (nB: median: 0.007, iqr: 0.004 – 0.015, B: median: 0.012, iqr: 0.006 – 0.023; p = 0.016, Wilcoxon rank-sum test).

Both bursting and non-bursting MCs preferentially fired during the trough of RR and theta oscillation (Figure 4E(a), F(a)) and the mean phase angle of locking did not differ for the two patterns of oscillatory activity (RR: nB: median: 0.917*π, iqr: 0.831*π – 0.968*π; B: median: 0.916*π, iqr: 0.761*π – 1.049*π; p= 0.803, circ_cmtest; theta: nB: median: 0.971*π, iqr: 0.925*π – 1.031*π; B: median: 0.948*π, iqr: 0.911*π – 1.006*π; p= 0.871, circ_cmtest). Reflecting the tighter coupling of bursting units to the theta band oscillations, the spike triggered average (STA) for the band-pass filtered LFP in theta range (4-12 Hz) showed higher absolute values for bursting units when compared to the non-bursting ones (Figure 4F(d); nB: median:1.077, iqr: 0.797 – 2.009; B: median: 2.000, iqr: 1.328 – 2.742; p = 0.031; Wilcoxon rank-sum test). In contrast, STA for RR was similar for the two types of units (Figure 4F(b), nB: median: 2.312, iqr: 1.299 – 4.773; B: median: 2.117, iqr: 1.141 – 3.294, p = 0.248; Wilcoxon rank-sum test). Moreover, the bursting and non-bursting units did not differ in their lag of the minimal STA deflection for both rhythms (RR: nB: median: 0.013, iqr: 0.009 – 0.025; B: median: 0.011, iqr: -0.005 – 0.027; p = 0.231; theta: nB: median: 0.002, iqr: -0.002 – 0.005 B: median: 0.002, iqr: -0.001 – 0.006; p = 0.808; Wilcoxon rank-sum test).

Taken together, these results indicate that bursting MCs tightly couple to theta oscillations in the neonatal OB.

### Mitral cells preferentially fire during theta events in the LEC

MCs have been found to directly project to limbic areas, such as LEC (Igarashi *et al.*, 2012). These projections emerge early in life and towards the end of the first postnatal week, actively contribute to the generation of theta band oscillatory events in the neonatal LEC (Gretenkord *et al.*, 2019) (Figure 5A). In line with other cortical areas, the LEC shows discontinuous large amplitude events with frequencies within 4-12 Hz range (theta events) that are superimposed on a continuous low-amplitude rhythm at 2-4 Hz (RR) (Figure 5B & C) (Hartung *et al.*, 2016; Gretenkord *et al.*, 2019). To distinguish how bursting and non-bursting MCs influence the entorhinal network activity, we examined the temporal relationship between the firing rate of bursting and non-bursting units in the neonatal OB and oscillatory activity in LEC. Both bursting and non-bursting MCs showed significantly higher normalized firing rates during entorhinal theta events when compared to time windows lacking those (theta events: nB: median: 1.183, iqr: 1.094 – 1.253; B: median: 1.315, iqr: 1.234 – 1.493; no events: nB: median: 0.820, iqr: 0.692 – 0.893; B: median: 0.640, iqr: 0.557 – 0.775; p(nB) = 3.79*10^−6^, p(B) = 1.17*10^−10^, Wilcoxon sign-rank test) (Figure 5B, D & E). However, the firing rate increase during theta events was more prominent for bursting than for non-bursting MCs (p = 1.69*10^4^, Wilcoxon rank-sum test). In contrast, bursting MCs had lower normalized firing rates during time windows lacking theta events (p = 8.68*10^−5^, Wilcoxon rank-sum test). These results demonstrate that among MCs the bursting ones have the tightest temporal correlation with the entorhinal theta oscillations.

**Figure 5:**
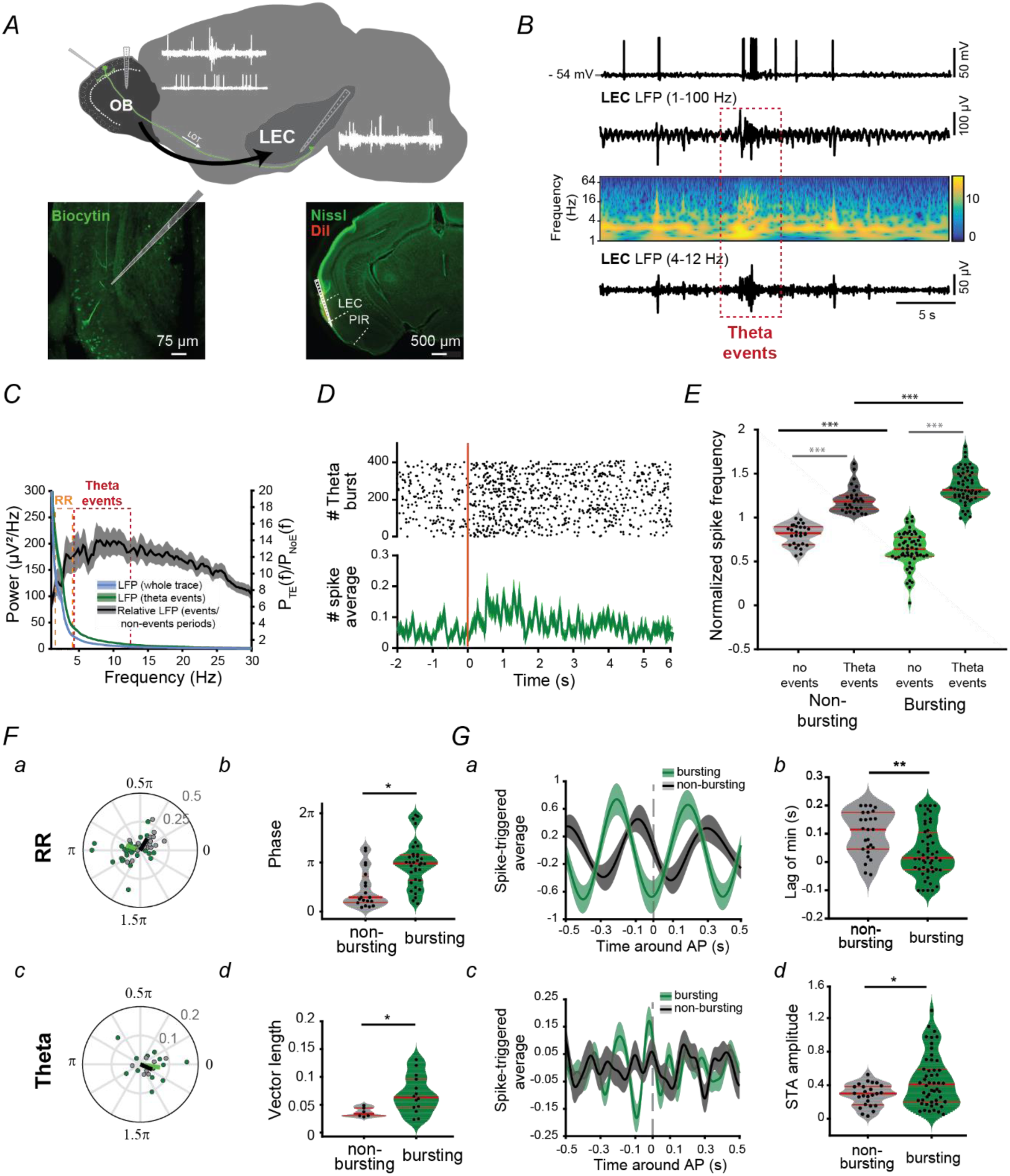
Bursting and non-bursting mitral cells have different phase locking dynamics to RR and theta events in the neonatal LEC. A, Top, schematic showing the recording configuration for simultaneous LFP and whole-cell patch-clamp recordings in the dorsal OB as well as for LFP recordings in LEC. LOT: lateral olfactory tract. Bottom, Confocal image displaying a biocytin-filled mitral cell (green) in the OB of a P8 mouse (left). Digital photomontage reconstructing the track of the multi-site DiI-labeled recording electrode (red) in a Nissl-stained coronal section of the LEC from a P9 mouse (right). PIR: piriform cortex B, Top, action potentials recorded simultaneously with the LFP from a MC at resting membrane potential in whole-cell current-clamp configuration. Bottom, extracellular LFP recording of the oscillatory activity in the LEC of a P9 mouse displayed after band-pass filtering (1-100 Hz, 4-12 Hz) and accompanied by the wavelet spectrogram at identical timescale. Note the temporal correlation between the discontinuous theta event (marked by red box) and action potentials. C, Power spectra (mean ± SEM) of LFP in LEC for whole recording (blue), during discontinuous bursts (green) as well as of theta bursts normalized to non-bursting activity (black). D, Representative raster plot (top) and averaged number of spikes (bottom) from an extracellularly recorded bursting unit aligned to the start time of each theta event in LEC (red line). E, Violin plot displaying the normalized firing rates of *in vivo* recorded non-bursting (gray) and bursting (blue) MCs during theta events in LEC (dark colors) and time windows lacking those (light colors). F, Phase locking of MC units to RR and theta events in LEC. Polar plots displaying the phase locking of significantly locked bursting (green) and non-bursting (gray) units to RR (a) and theta (c). Violin plots showing the phase for bursting (green) and non-bursting units phase locked to RR (b) and the mean resulting vector length for bursting (green) and non-bursting units phase locked to theta (d). Only significantly locked units where used for analysis. G, Spike triggered average for RR (a) and theta (c) in LEC around the time of action potential occurrence for bursting (green) and non-bursting (grey) MCs. Corresponding violin plots showing the lag of the minimal deflection of the STA for RR (b) and the absolute STA amplitude around the time of action potential occurrence for theta (d). (*p<0.05, **p<0.01, *** p < 0.001; Wilcoxon rank-sum test (black), Wilcoxon sign-rank test (gray) or circ_cmtest (matlab) for phase values). For violin plots the red lines correspond to median (thick line) as well as 1. and 3. quantiles of the distribution (thin lines).

To deepen the investigation of the temporal relationship between MCs in OB and network activity of LEC, we assessed the phase locking of bursting and non-bursting units extracellularly recorded in the MC layer to the entorhinal theta and RR. MCs were phase locked to the continuous RR (non-bursting: 73 %, bursting 69%) and to a lower extent, to theta events (non-bursting: 27%, bursting 27%). The locking strength to RR was comparable for significantly locked bursting and non-bursting MCs (nB: median: 0.109. iqr: 0.069 – 0.149 B: median: 0.104, iqr: 0.055 – 0.139; p = 0.578; Wilcoxon rank-sum test). However, bursting MCs were locked at a different phase (through) than non-bursting ones (peak) (Figure 5F(b); nB: median: 0.288*π, iqr: 0.184* π – 0.716*π; B: median: 0.980* π, iqr: 0.639* π – 1.154* π; p= 0.002, circ_cmtest). The strength of locking to theta phase in LEC was higher for bursting (Figure 5F(d); B: median: 0.063, iqr: 0.045 - 0.096) when compared to non-bursting MCs (nB: median: 0.034, iqr: 0.030 - 0.044; p= 0.035; Wilcoxon rank-sum test). The mean phase angle of locking did not differ for the two firing patterns (nB: median: 1.051*π, iqr: 0.122*π – 1.693*π; B: median: 1.278*π, iqr: 0.278*π – 1.803*π; p = 0.537, circ_cmtest). Confirming previously described results, STA for RR in LEC was similar for bursting and non-bursting MCs (RR: nB: median: 0.645, iqr: 0.494-1.295; B: median: 0.710, iqr: 0.338 - 1.474; p = 0.720, Wilcoxon rank-sum test). In contrast, STA for theta phase showed higher absolute values for bursting units when compared to the non-bursting ones (Figure 5G(d); theta: nB: median: 0.300, iqr: 0.165 - 0.384; B: median: 0.408, iqr: 0.202 - -0.591; p = 0.036, Wilcoxon rank-sum test). The two populations of MCs showed differences in their lag of the minimal STA deflection for entorhinal RR (Figure 5G(b); nB: median: 0.115, iqr: 0.046 – 0.176; B: median: 0.015, iqr: -0.027 – 0.104; p = 0.003, Wilcoxon rank-sum test) but not theta phase (nB: median: -0.039, iqr: -0.080 – 0.009; B: median: -0.050, iqr: -0.093 – 0.022; p = 0.354, Wilcoxon rank-sum test).

Taken together these data show that among MCs, the bursting cells have the strongest temporal relationship with the theta events in LEC. This might suggest that the bursting MCs preferentially relay OB information to LEC and, by these means, facilitates the oscillatory entrainment of limbic circuits.

## Discussion

The processing ability of the mature brain relies on the temporal coupling of neuronal firing to network oscillations and the resulting entrainment of local and large-scale networks. Already during early development, neurons in sensory and limbic brain areas generate distinct firing patterns and assemble into circuits synchronized in different frequency bands (Khazipov *et al.*, 2004; Brockmann *et al.*, 2011; Minlebaev *et al.*, 2011; Bitzenhofer *et al.*, 2015; Hartung *et al.*, 2016; Ahlbeck *et al.*, 2018). At this age, when most sensory inputs do not reach the cortex, spontaneous and stimulus-induced activation of the LEC via direct axonal projections of MCs might boost the development of the limbic circuitry (Gretenkord *et al.*, 2019). The mechanisms of communication between OB and LEC are still largely unresolved. One critical question is how the firing of MCs entrains entorhinal circuits. Combining *in vivo* and *in vitro* patch-clamp recordings from MCs with extracellular recordings of LFP and spiking activity in OB and LEC, we demonstrate that (i) during early postnatal development (P8 – 10) MCs display either irregular busting or non-bursting firing characteristics, (ii) irregular bursting MCs display less pronounced hyperpolarization-evoked sag currents when compared to non-bursting MCs, (iii) the firing of bursting MCs concentrate during theta events and is more precisely locked to the phase of theta oscillations in the neonatal OB, and (iv) bursting MCs temporally relate to oscillatory patterns in neonatal LEC. These findings suggest that predominantly bursting MCs synchronize neuronal ensembles in discontinuous theta oscillations in the developing OB and LEC.

While the intrinsic firing properties of MCs have been investigated extensively in adult rodents (Yu *et al.*, 1993; Balu *et al.*, 2004; Nica *et al.*, 2010; Angelo & Margrie, 2011), little is known about their firing properties during early postnatal development. To our knowledge, only one study has previously investigated the passive and active properties of MCs during the first two postnatal weeks *in vitro.* It reports an age-dependent decrease in membrane time constant and input resistance but no change in membrane capacity or resting membrane potential (Yu *et al.*, 2015). The study classified MCs as either bursting (stuttering) or non-bursting according to their firing patterns. Here, we extended these findings by demonstrating that also *in vivo* MCs show bursting or non-bursting firing patterns already at neonatal age. These firing patterns were mostly independent from the recording condition, passive membrane properties as well as from the firing rates of the recorded neurons. Bursting MCs show less pronounced sag currents compared to non-bursting MCs, indicating lower expression of hyperpolarization-activated cation (HCN) channels and therefore h-currents (Robinson & Siegelbaum, 2003; Angelo & Margrie, 2011). This is well in line with several studies in adult rodents (Angelo & Margrie, 2011; Burton & Urban, 2014) as well as *in vitro* studies in neonatal mice (Yu *et al.*, 2015) which demonstrate a link between sag potential amplitude and firing regularity of mitral cells.

Sag amplitudes of MCs were shown to decrease significantly over the first postnatal weeks leading to more irregular firing characteristics and therefore more stuttering firing pattern of MCs (Yu *et al.*, 2015). These changes together with changes in intrinsic properties of MCs suggests that MCs attune to faster network oscillations across development. While MCs during early postnatal development attune preferentially to theta oscillations, mature MCs are more likely to attune to gamma oscillations (Yu *et al.*, 2015). How the different firing pattern of MCs emerge is yet not fully understood. Yu et al. suggest that bursting MCs show a lower level of HCN channels. Moreover, spike clustering in mature MCs has been linked to I_D_-like potassium currents and subthreshold TTX-sensitive sodium currents (Balu *et al.*, 2004). However, the spontaneous firing patter of MCs reflect most likely a combination from intrinsic properties and local network interactions (Leng *et al.*, 2014). Accordingly, we show a slightly lower portion of bursting MCs *in vitro* compared to our *in vivo* conditions. This might be due to truncated centrifugal afferents from other brain areas in *in vitro* preparations. Centrifugal feedback as well as neuromodulation might critically shape MC firing pattern *in vivo*. Another factor that might impact neuronal firing is anesthesia. However, we have shown recently that while urethan dampens the overall level of activity it does not change the oscillatory events per se and the temporally correlated firing within developing OB-LEC networks (Chini *et al.*, 2019; Gretenkord *et al.*, 2019). Finally, it is likely that ongoing sensory sampling plays an important role in shaping the firing pattern of MCs. Further research is needed to reveal the developmental mechanisms that shape the patterning of mitral cell firing during passive breathing as well as during active odor sampling.

The present study reveals that the temporal coupling between MC firing and theta events is more prominent for bursting than for non-bursting MCs. These findings suggest a stronger contribution of bursting MCs when compared to non-bursting MCs to the generation of theta events in the neonatal OB. Intrinsically bursting neurons have been proposed as key elements that entrain developing circuits in oscillatory rhythms (Egorov & Draguhn, 2013). For example, bursting neurons have been shown to contribute to early network oscillations in the entorhinal cortex (Sheroziya *et al.*, 2009). Moreover, it has been suggested that spontaneously bursting starburst cells boost the generation of retinal waves (Zheng *et al.*, 2006) and intrinsic bursting CA3 neurons shape hippocampal giant depolarizing potentials (GDPs) (Sipilä *et al.*, 2006). Additionally, in the mature brain, bursting firing might set functional networks to preferred frequencies. For example, mature bursting MCs attune preferentially to the theta rhythm (Balu *et al.*, 2004). Bursting MCs in the developing OB might thus similarly act as pacemakers, shaping the neonatal theta rhythm.

An important open question is how OB activity shapes the maturation of the OB-LEC circuitry during neonatal development. The present data suggest that bursting MCs preferentially relay activity from the OB to LEC during neonatal development via axonal projections of MCs that target LEC already around birth (Gretenkord *et al.*, 2019). Bursting neurons have been proposed to transmit information more reliable compared to single spikes (Lisman, 1997). Therefore, bursting MCs might efficiently activate entorhinal layer II/III neurons that have been shown to have a critical role for the generation of theta oscillations (Alonso & Llinás, 1989; Alonso & Klink, 1993). Additionally, rhythmically firing layer II/III neurons in the medial entorhinal cortex preferentially fire during the negative phase of the theta rhythm (4-6 Hz), while non-rhythmic cells fire during the positive phase (Alonso & Garcia-Austt, 1987). In line with this notion, we also report that bursting MCs show a different phase relationship to RR (2-4 Hz) in LEC compared to non-bursting MCs. Moreover, bursting activity has been related to plasticity mechanism (Pike *et al.*, 1999) in the HP. Based on studies investigating the synaptic refinement of retinal ganglion cell in the developing visual system, it has been proposed that during development, burst-time dependent plasticity rather than spike-time dependent plasticity plays an important role in the strengthening of synapses (Butts *et al.*, 2007; Gjorgjieva *et al.*, 2009; Butts & Kanold, 2010). Therefore, it is tempting to speculate that bursting MCs strengthen the connectivity between OB and LEC during neonatal development, thereby contributing to the maturation of the network.

In sum, we propose that bursting MCs act as pacemakers for theta events in OB and reliably activate entorhinal networks at neonatal age, thereby facilitating the oscillatory entrainment of developing limbic circuits.

## Acknowledgments

We thank Drs. Marc Spehr and Yoram Ben-Shaul for providing the analytical tools for burst detection, A. Marquardt, A. Dahlmann, P. Putthoff and K. Titze for excellent technical assistance.

## Additional Information

### Data Availability

The data that supports the findings of this study are available in the supplementary material of this article.

### Author Contributions

I.L.H.-O. conceived the study and designed the experiments. J.K.K and S.G. carried out the experiments. J.K.K analyzed the data. J.K.K, and I.L.H.-O. interpreted the data. J.K.K. and I.L.H.-O. wrote the article. All authors discussed and commented on the manuscript.

## References

Adam Y & Mizrahi A. (2011). Long-term imaging reveals dynamic changes in the neuronal composition of the glomerular layer. Journal of Neuroscience 31, 7967–7973.

Ahlbeck J, Song L, Chini M, Bitzenhofer SH & Hanganu-Opatz IL. (2018). Glutamatergic drive along the septo-temporal axis of hippocampus boosts prelimbic oscillations in the neonatal mouse. Elife 7, e33158.

Alonso A & Garcia-Austt E. (1987). Neuronal sources of theta rhythm in the entorhinal cortex of the rat. Experimental brain research 67, 493–501.

Alonso A & Klink R. (1993). Differential electroresponsiveness of stellate and pyramidal-like cells of medial entorhinal cortex layer II. Journal of neurophysiology 70, 128–143.

Alonso A & Llinás RR. (1989). Subthreshold Na+-dependent theta-like rhythmicity in stellate cells of entorhinal cortex layer II. Nature 342, 175–177.

Angelo K & Margrie TW. (2011). Population diversity and function of hyperpolarization-activated current in olfactory bulb mitral cells. Scientific reports 1, 50.

Angelo K, Rancz EA, Pimentel D, Hundahl C, Hannibal J, Fleischmann A, Pichler B & Margrie TW. (2012). A biophysical signature of network affiliation and sensory processing in mitral cells. Nature 488, 375–378.

Balu R, Larimer P & Strowbridge BW. (2004). Phasic stimuli evoke precisely timed spikes in intermittently discharging mitral cells. Journal of neurophysiology 92, 743–753.

Barbour B. (2011). Electronics for electrophysiologists.

Bitzenhofer SH, Ahlbeck J, Wolff A, Wiegert JS, Gee CE, Oertner TG & Hanganu-Opatz IL. (2017). Layer-specific optogenetic activation of pyramidal neurons causes beta–gamma entrainment of neonatal networks. Nature communications 8, 14563.

Bitzenhofer SH, Sieben K, Siebert KD, Spehr M & Hanganu-Opatz IL. (2015). Oscillatory activity in developing prefrontal networks results from theta-gamma-modulated synaptic inputs. Cell reports 11, 486–497.

Blanchart A, De Carlos JA & López-Mascaraque L. (2006). Time frame of mitral cell development in the mice olfactory bulb. Journal of Comparative Neurology 496, 529–543.

Brockmann MD, Pöschel B, Cichon N & Hanganu-Opatz IL. (2011). Coupled oscillations mediate directed interactions between prefrontal cortex and hippocampus of the neonatal rat. Neuron 71, 332–347.

Burton SD & Urban NN. (2014). Greater excitability and firing irregularity of tufted cells underlies distinct afferent-evoked activity of olfactory bulb mitral and tufted cells. The Journal of physiology 592, 2097–2118.

Butts DA & Kanold PO. (2010). The applicability of spike time dependent plasticity to development. Frontiers in synaptic neuroscience 2, 30.

Butts DA, Kanold PO & Shatz CJ. (2007). A burst-based “Hebbian” learning rule at retinogeniculate synapses links retinal waves to activity-dependent refinement. PLoS biology 5.

Chini M, Gretenkord S, Kostka JK, Pöpplau JA, Cornelissen L, Berde CB, Hanganu-Opatz IL & Bitzenhofer SH. (2019). Neural correlates of anesthesia in newborn mice and humans. Frontiers in neural circuits 13, 38.

Cichon NB, Denker M, Grün S & Hanganu-Opatz IL. (2014). Unsupervised classification of neocortical activity patterns in neonatal and pre-juvenile rodents. Frontiers in neural circuits 8, 50.

Egorov AV & Draguhn A. (2013). Development of coherent neuronal activity patterns in mammalian cortical networks: common principles and local hetereogeneity. Mechanisms of development 130, 412–423.

Gjorgjieva J, Toyoizumi T & Eglen SJ. (2009). Burst-time-dependent plasticity robustly guides ON/OFF segregation in the lateral geniculate nucleus. PLoS computational biology 5.

Gorin M, Tsitoura C, Kahan A, Watznauer K, Drose DR, Arts M, Mathar R, O’Connor S, Hanganu-Opatz IL & Ben-Shaul Y. (2016). Interdependent conductances drive infraslow intrinsic rhythmogenesis in a subset of accessory olfactory bulb projection neurons. Journal of Neuroscience 36, 3127–3144.

Gretenkord S, Kostka JK, Hartung H, Watznauer K, Fleck D, Minier-Toribio A, Spehr M & Hanganu-Opatz IL. (2019). Coordinated electrical activity in the olfactory bulb gates the oscillatory entrainment of entorhinal networks in neonatal mice. PLoS biology 17, e2006994.

Hartung H, Brockmann MD, Pöschel B, De Feo V & Hanganu-Opatz IL. (2016). Thalamic and entorhinal network activity differently modulates the functional development of prefrontal–hippocampal interactions. Journal of Neuroscience 36, 3676–3690.

Hinds JW & Hinds PL. (1976). Synapse formation in the mouse olfactory bulb. II. Morphogenesis. Journal of Comparative Neurology 169, 41–61.

Hirata T, Shioi G, Abe T, Kiyonari H, Kato S, Kobayashi K, Mori K & Kawasaki T. (2019). A Novel Birthdate-Labeling Method Reveals Segregated Parallel Projections of Mitral and External Tufted Cells in the Main Olfactory System. eNeuro 6.

Igarashi KM, Ieki N, An M, Yamaguchi Y, Nagayama S, Kobayakawa K, Kobayakawa R, Tanifuji M, Sakano H & Chen WR. (2012). Parallel mitral and tufted cell pathways route distinct odor information to different targets in the olfactory cortex. Journal of Neuroscience 32, 7970–7985.

Khazipov R, Sirota A, Leinekugel X, Holmes GL, Ben-Ari Y & Buzsáki G. (2004). Early motor activity drives spindle bursts in the developing somatosensory cortex. Nature 432, 758–761.

Kollo M, Schmaltz A, Abdelhamid M, Fukunaga I & Schaefer AT. (2014). ’Silent’mitral cells dominate odor responses in the olfactory bulb of awake mice. Nature neuroscience 17, 1313.

Leng G, Hashimoto H, Tsuji C, Sabatier N & Ludwig M. (2014). Discharge patterning in rat olfactory bulb mitral cells in vivo. Physiological reports 2, e12021.

Lin DM, Wang F, Lowe G, Gold GH, Axel R, Ngai J & Brunet L. (2000). Formation of precise connections in the olfactory bulb occurs in the absence of odorant-evoked neuronal activity. Neuron 26, 69–80.

Lisman JE. (1997). Bursts as a unit of neural information: making unreliable synapses reliable. Trends in neurosciences 20, 38–43.

Logan DW, Brunet LJ, Webb WR, Cutforth T, Ngai J & Stowers L. (2012). Learned recognition of maternal signature odors mediates the first suckling episode in mice. Current biology 22, 1998–2007.

Luskin MB & Price JL. (1983). The topographic organization of associational fibers of the olfactory system in the rat, including centrifugal fibers to the olfactory bulb. Journal of comparative neurology 216, 264–291.

Minlebaev M, Colonnese M, Tsintsadze T, Sirota A & Khazipov R. (2011). Early gamma oscillations synchronize developing thalamus and cortex. Science 334, 226–229.

Monier C, Fournier J & Frégnac Y. (2008). In vitro and in vivo measures of evoked excitatory and inhibitory conductance dynamics in sensory cortices. Journal of neuroscience methods 169, 323–365.

Nica R, Matter SF & Griff ER. (2010). Physiological evidence for two classes of mitral cells in the rat olfactory bulb. Brain research 1358, 81–88.

Padmanabhan K & Urban NN. (2010). Intrinsic biophysical diversity decorrelates neuronal firing while increasing information content. Nature neuroscience 13, 1276.

Pike FG, Meredith RM, Olding AW & Paulsen O. (1999). Postsynaptic bursting is essential for ‘Hebbian’induction of associative long-term potentiation at excitatory synapses in rat hippocampus. The Journal of physiology 518, 571–576.

Robinson RB & Siegelbaum SA. (2003). Hyperpolarization-activated cation currents: from molecules to physiological function. Annual review of physiology 65, 453–480.

Rossant C, Kadir SN, Goodman DF, Schulman J, Hunter ML, Saleem AB, Grosmark A, Belluscio M, Denfield GH & Ecker AS. (2016). Spike sorting for large, dense electrode arrays. Nature neuroscience 19, 634.

Sheroziya MG, und Halbach OvB, Unsicker K & Egorov AV. (2009). Spontaneous bursting activity in the developing entorhinal cortex. Journal of Neuroscience 29, 12131–12144.

Shinomoto S. (2010). Estimating the firing rate. In Analysis of Parallel Spike Trains, pp. 21–35. Springer.

Siapas AG, Lubenov EV & Wilson MA. (2005). Prefrontal phase locking to hippocampal theta oscillations. Neuron 46, 141–151.

Sipilä ST, Huttu K, Voipio J & Kaila K. (2006). Intrinsic bursting of immature CA3 pyramidal neurons and consequent giant depolarizing potentials are driven by a persistent Na+ current and terminated by a slow Ca2+-activated K+ current. European Journal of Neuroscience 23, 2330–2338.

Suter BA, O’Connor T, Iyer V, Petreanu L, Hooks BM, Kiritani T, Svoboda K & Shepherd GM. (2010). Ephus: multipurpose data acquisition software for neuroscience experiments. Frontiers in neural circuits 4, 100.

Teicher MH & Blass EM. (1977). First suckling response of the newborn albino rat: the roles of olfaction and amniotic fluid. Science 198, 635–636.

Vinck M, van Wingerden M, Womelsdorf T, Fries P & Pennartz CM. (2010). The pairwise phase consistency: a bias-free measure of rhythmic neuronal synchronization. Neuroimage 51, 112–122.

Walz A, Omura M & Mombaerts P. (2006). Development and topography of the lateral olfactory tract in the mouse: imaging by genetically encoded and injected fluorescent markers. Journal of neurobiology 66, 835–846.

Welker W. (1964). Analysis of sniffing of the albino rat 1. Behaviour 22, 223–244.

Witter MP. (2007). The perforant path: projections from the entorhinal cortex to the dentate gyrus. Progress in brain research 163, 43–61.

Wouterlood F, Mugnaini E & Nederlof J. (1985). Projection of olfactory bulb efferents to layer I GABAergic neurons in the entorhinal area. Combination of anterograde degeneration and immunoelectron microscopy in rat. Brain research 343, 283–296.

Yu G-Z, Kaba H, Saito H & Seto K. (1993). Heterogeneous characteristics of mitral cells in the rat olfactory bulb. Brain research bulletin 31, 701–706.

Yu Y, Burton SD, Tripathy SJ & Urban NN. (2015). Postnatal development attunes olfactory bulb mitral cells to high-frequency signaling. Journal of neurophysiology 114, 2830–2842.

Zheng J, Lee S & Zhou ZJ. (2006). A transient network of intrinsically bursting starburst cells underlies the generation of retinal waves. Nature neuroscience 9, 363–371.

